# Exploring epitope and functional diversity of anti-SARS-CoV2 antibodies using AI-based methods

**DOI:** 10.1101/2020.12.23.424199

**Authors:** C. Dumet, Y. Jullian, A. Musnier, Ph. Rivière, N. Poirier, H. Watier, T. Bourquard, A. Poupon

## Abstract

Since the beginning of the COVID19 pandemics, an unprecedented research effort has been conducted to analyze the antibody responses in patients, and many trials based on passive immunotherapy — notably monoclonal antibodies — are ongoing. Twenty-one antibodies have entered clinical trials, 6 having reached phase 2/3, phase 3 or having received emergency authorization. These represent only the tip of the iceberg, since many more antibodies have been discovered and represent opportunities either for diagnosis purposes or as drug candidates. The main problem facing laboratories willing to develop such antibodies is the huge task of analyzing them and choosing the best candidate for exhaustive experimental validation. In this work we show how artificial intelligence-based methods can help in analyzing large sets of antibodies in order to determine in a few hours the best candidates in few hours. The MAbCluster method, which only requires knowledge of the amino acid sequences of the antibodies, allows to group the antibodies having the same epitope, considering only their amino acid sequences and their 3D structures (actual or predicted), and to infer some of their functional properties. We then use MAbTope to predict the epitopes for all antibodies for which they are not already known. This allows an exhaustive comparison of the available epitopes, but also gives a synthetic view of the possible combinations. Finally, we show how these results can be used to predict which antibodies might be affected by the different mutations arising in the circulating strains of the virus, such as the N501Y mutation that has started to spread in Great-Britain.

---

Faced with the gravity and spread of the COVID19 pandemics, pharmaceutical companies, biotechs and hospitals have launched therapeutic trials, with various degree of control^*,1^. Notably, many anti-viral, anti-malaria and anti-inflammatory molecules, initially developed for different applications, have been tested, with very limited success^2^.

There is now ample evidence that the main pathway allowing the SARS-CoV-2 virus to infect human cells relies on the binding of the receptor binding domain (RBD) of the spike protein to the human angiotensin-converting enzyme 2 (ACE2)^3^. Other human proteins were shown to interact with the spike protein of the SARS-CoV2, and suspected to provide alternative routes for the entry of the virus in the human cells. For example, basigin was first thought to be such an alternative route. However, although the interaction could be demonstrated experimentally^4^, its role in infection could not be proven^5^. It was also shown that the spike protein, through its multiple glycosylations, interacts with multiple immune receptors^6^, which might represent alternative internalization pathways.

The research effort has therefore concentrated on blocking the RBD-ACE2 interaction. Significant progress has been made using engineered ACE2 versions that could be injected in order to trap the virus^7^, and trials are ongoing^†^. The real successes have been obtained using monoclonal antibodies binding the RBD, therefore preventing the entry of the virus in the host’s cells. To date, twenty-one antibodies targeting SARS-CoV-2, all binding to the receptor binding domain (RBD) of the spike protein, have entered therapeutic trials (Supp. table 1), 6 having reached phase 2/3, phase 3 or having received emergency authorization^8^.

## Therapeutic candidates epitope comparison

For only 8 of the monoclonal antibodies under trial has the amino acid sequence been released. For 2 of them the 3D structure of the complex between the antibody and the RBD is known. For the 6 others the epitope was determined using MAbTope^9^ (Figure 1).

**Figure 1:**
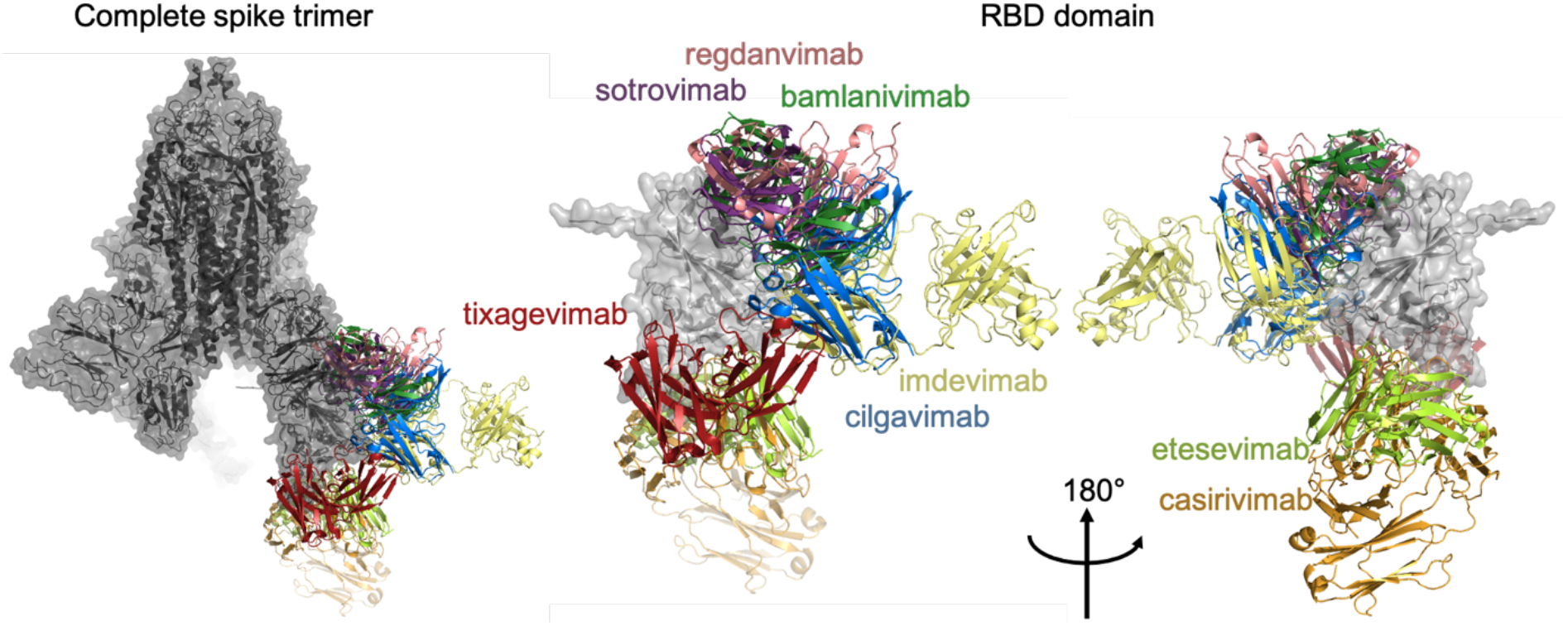
8 therapeutic candidates in complex with the spike protein.

As expected, the pairs of antibodies that are being tested in combination and have reached phase 3 (tixagevimab + cilgavimab) or are already authorized (etevesimab + bamlanivimab and casirivimab + imdevimab), correspond to non-overlapping epitopes. This analysis also shows which other combinations might represent new therapeutic opportunities, and which ones would probably be ineffective because the antibodies bind the same epitopes.

Surprisingly, sotrovimab^10^ and imdevimab^11^ bind to epitopes that do not overlap with the region interacting with ACE2. The 3D structure of the complex between RBD, ACE2 and B^0^AT1 (PDB: 6M17^12^) gives good hypotheses. Indeed, the authors show that ACE2 forms dimers, and binding of these antibodies to the RBD prevents the binding of the RBD on ACE2, not directly, but through steric clashes with the second ACE2 monomer (Supp. figure 1). Thus, the complexes spike-sotrovimab and spike-imdevimab cannot bind the ACE2 dimer, which seems to be its functional state, and are therefore neutralizing the virus entry.

## Clustering antibodies to predict functional properties

We conducted an exhaustive patent and literature review of the antibodies targeting the SARS-CoV-2 spike protein, but also the spike proteins of other coronaviruses, in particular SARS-CoV-1 and MERS-CoV. We found 1569 antibodies for which the sequences were available. We used MAbCluster to group these antibodies by epitopes (Figure 2). The principle of this AI-based method is to measure the similarity between the antibodies, based on both the sequence and the structure of the complementary determining regions (CDRs)^13^. We have already demonstrated that two antibodies having sufficient similarity (as measured by our algorithm) bind to the same epitope(s) of the same target (or targets). For clarity, only clusters having more than 6 members (33 clusters out of 703, accounting for 561 antibodies out of 1571) are highlighted in Figure 2.

**Figure 2:**
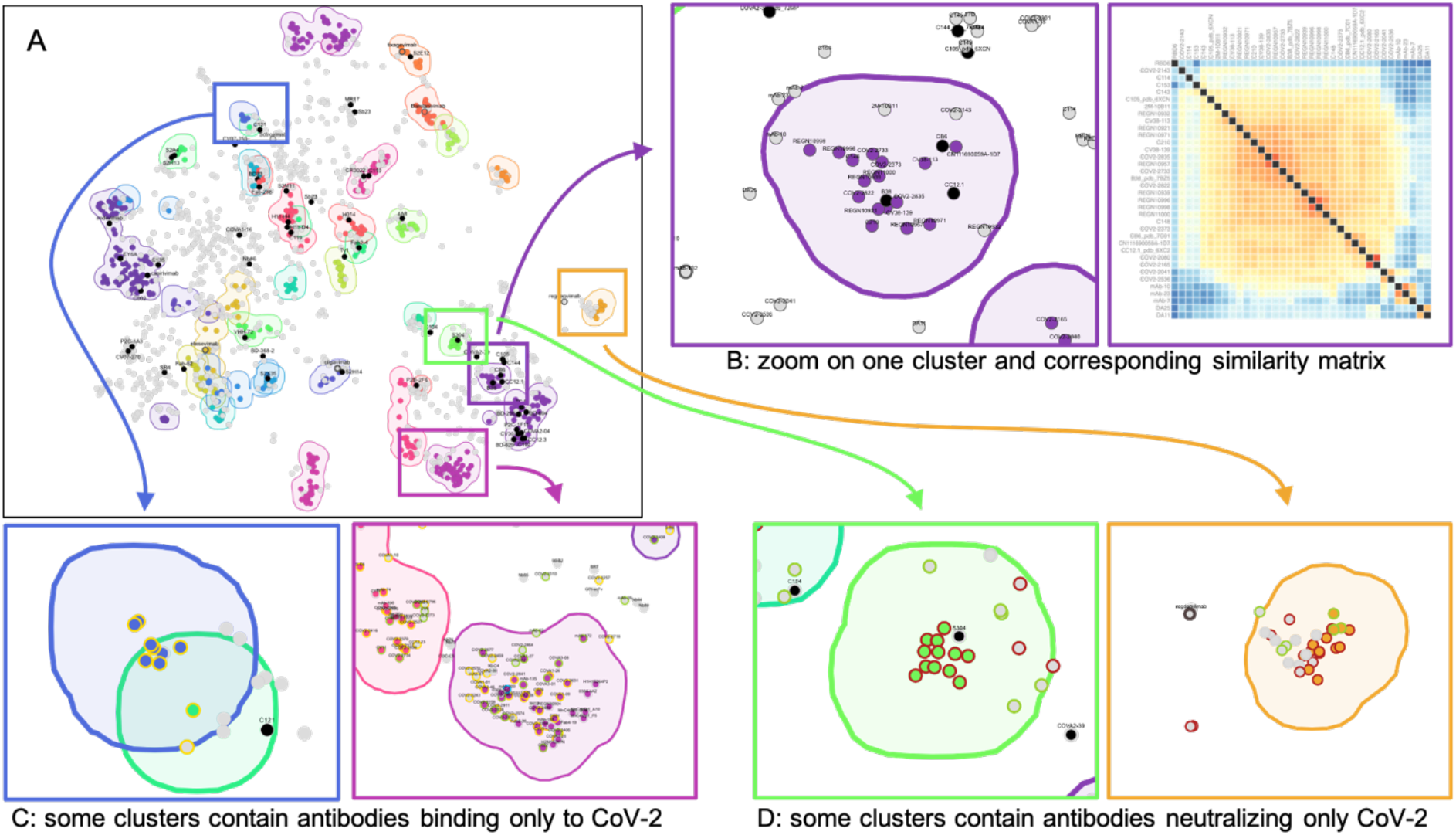
A- UMAP view of the clustering of the 1569 antibodies. Dots representing antibodies whose epitopes are known are indicated in black, clusters with 7 or more members, and the antibodies belonging to them are indicated in various colors. Grey dots represent antibodies belonging to smaller clusters. B- zoom on a region of the map and corresponding similarity matrix of the antibodies within the zoomed region. C- Zoom on two clusters, antibodies binding only to CoV-2 (and not CoV-1) are represented as yellow circles, antibodies binding to both as green circles, antibodies binding only to CoV-1 (and not CoV-2) as blue circles, antibodies for which the information is not available as grey circles. D: zoom on two clusters, antibodies neutralizing CoV-2 are indicated as green circles, antibodies not neutralizing CoV-2 as red circles, antibodies for which the information is not available as grey circles.

This clustering can be used to very rapidly determine the epitopes of antibodies that are similar to an antibody whose epitope is known. For example, among the dots present in the zoomed region (Figure 2B), the corresponding similarity matrix allows to predict that all the antibodies of the cluster (those colored in violet) bind to the same epitope. This cluster contains COVA2-04, P2C-1F11, 7CJF, CV30, CC12.3, BD-629 and BD-604, for which the 3D structure of the complex with RBD have been determined, and which all have very similar epitopes, in the same region of the RBD. Thus, we can predict with good probability that all the antibodies of the cluster have epitopes corresponding to this same region.

This method can be used for very large numbers of antibodies, such as the sequencing of the antibodies remaining after bio-panning selection, and allows to rapidly select the most promising ones. For example, panel C of Figure 2 shows that although some clusters gather antibodies having different specificities (towards CoV-1 and/or CoV-2, right), others gather only antibodies having the same specificities, such as the cluster shown on the right in Figure 2C, which contains only antibodies binding to CoV-2 and not to CoV-1. Consequently, when analyzing a new antibody which falls within this cluster, one can reasonably make the assumption that this new antibody only binds CoV-2. A similar analysis can be made on the neutralizing capacities of antibodies. Figure 2D shows a zoom on two clusters of CoV-2-binding antibodies. Whereas the cluster on the right gathers both neutralizing and non-neutralizing antibodies, the cluster on the left only gathers non-neutralizing antibodies. Thus, an antibody falling in the latter cluster can be predicted as non-neutralizing.

## Insights from epitope mapping

By searching in the protein data bank, we found 48 additional anti-spike antibodies targeting the RBD, and for which the 3D structure of the complex is known. For all the remaining antibodies amongst the 1569, epitopes were predicted using MAbTope.

For visualization, we show only one representative of each cluster (having more than 6 members). The criteria used to choose the representative for each of these clusters was: (1) antibody for which the structure of the complex with RBD in known; (2) if none available, antibody for which the highest amount of information is known (binding to CoV-1, neutralization of CoV-2, neutralization of CoV-1, etc.). We also added to the visualization other antibodies for which the antibody-target 3D structure is known, but which do not belong to a cluster having more than 6 members (Supp. table 2). We then compute the overlap between the epitopes. The result is shown as a matrix on Figure 3. Details of the epitopes of the antibodies are given Supp. figure 2.

**Figure 3:**
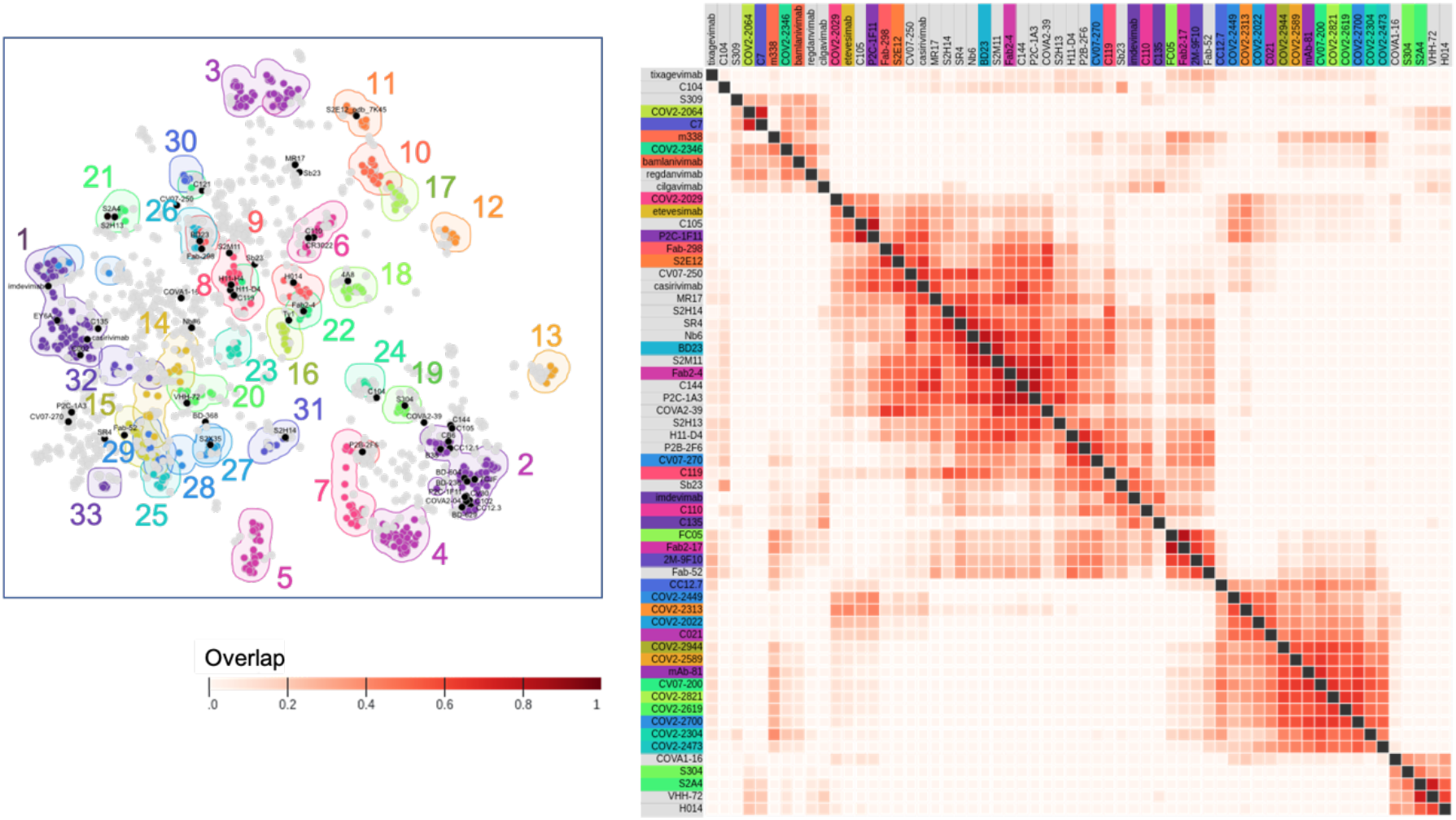
Epitope overlap. For each of the 33 clusters having more than 6 members (left), one representative was chosen (Table S1). For representatives with unknown antibody-target 3D structure, the epitope was determined using MAbTope. Antibodies with known antibody-target 3D structure not belonging to a cluster having more than 6 members were also considered. The pairwise overlaps between epitopes are plotted on the matrix (right).

Knowledge of the epitopes allows further interpretations of the conclusions inferred from the clustering. For example, epitope mapping of antibodies belonging to cluster #30 (Figure 2C, left) shows that these antibodies bind to a region of SARS-CoV-2 that is very different from the corresponding region in SARS-CoV-1, explaining why antibodies belonging to this cluster bind to SARS-CoV-2 and not SARS-CoV-1. On the contrary, cluster #4 (Figure 2C right) gathers antibodies whose epitopes contain both conserved and non-conserved regions. It is thus impossible to predict if antibodies within this cluster bind to both species or only one. Similarly, antibodies belonging to cluster #24 (Figure 2D left) bind to a region which presents important structural differences between the two viruses, and are neutralizing only to SARS-CoV-2. On the contrary, antibodies belonging to cluster #13 (Figure 2D right) bind to epitopes spanning on both conserved and non-conserved regions, and some of them are neutralizing either to one of the viruses or to both.

Comparison of the epitopes of all the antibodies studied here led us to define 10 binding regions. Given the shoe-like shape of the RBD, we named these regions as indicated in Figure 4.Unsurprisingly, none of the antibodies bind the cuff region, which corresponds to the N and C-term of the RBD, and is thus buried by the remaining of the spike. More surprising, no antibody binds the toe region, also it is well exposed in both opened and closed conformations.

**Figure 4:**
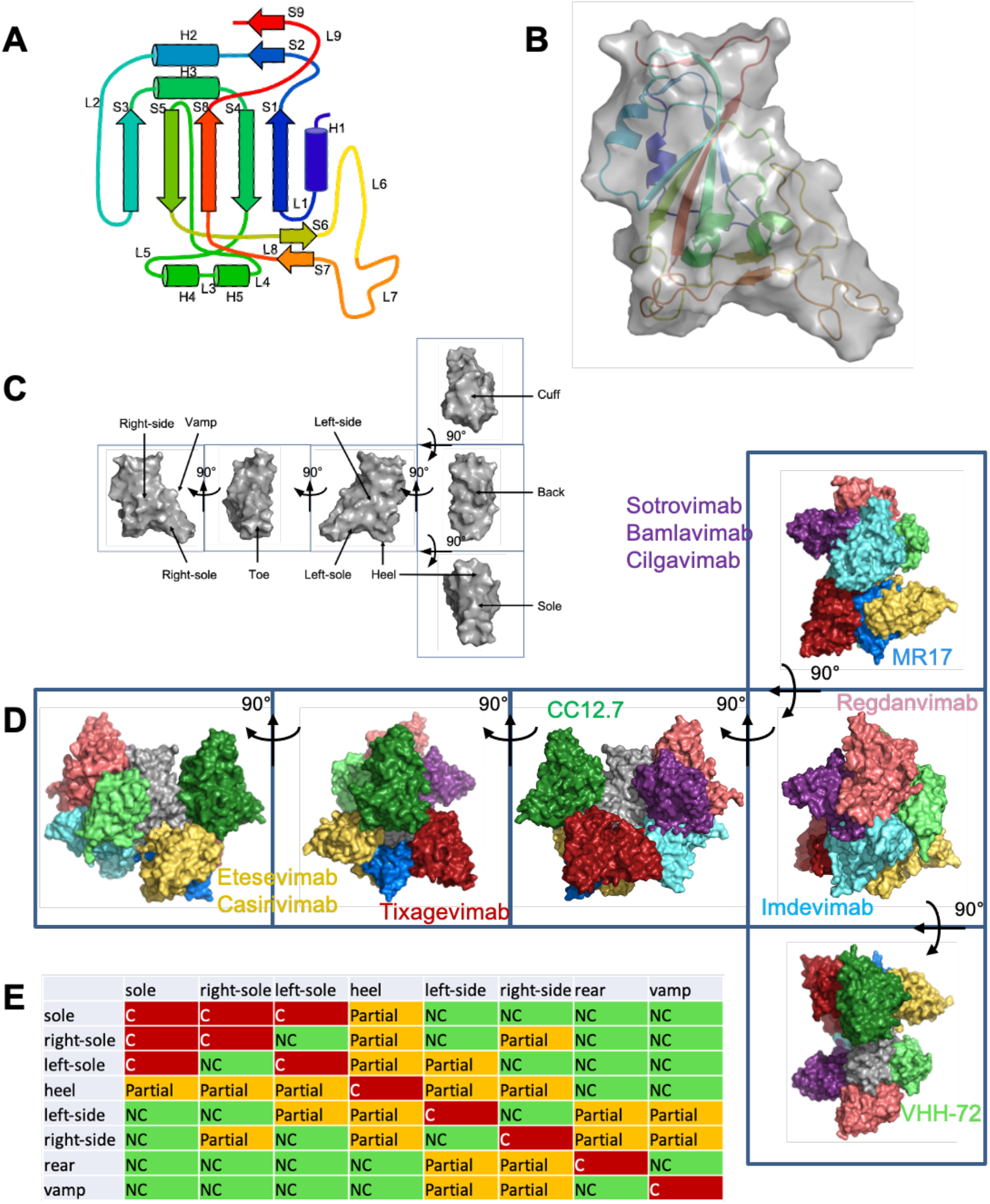
Defining binding regions of the RBD. A: secondary structure; B: surface representation; C: naming of the different regions. D: Binding of antibodies belonging to the 8 defined groups on the RBD. D: Global competition table between the antibodies belonging to the different regions, NC (green): non competing, Partial (Orange): partial competition (meaning that some antibodies of the first group compete with some antibodies of the second group); C (red): competing.

As can be seen from Table 1, the region to which the highest number of antibodies (among which etevesimab and casirivimab) bind is the right-sole, which is not surprising since this is the region interaction with ACE2. It is thus also unsurprising that a high proportion of these antibodies are neutralizing (95 out of 159 tested). This region also appears as specific to CoV-2, since the proportion of antibodies also binding CoV-1 is limited (48 out of 128 tested), mostly due to the fact that secondary structures L6, L7, S7 and S8 are very different in the two strains (Supp. figure 2). Many antibodies are binding to the sole region (125). Although antibodies binding to this region should also interfere with the binding to ACE2, the proportion of neutralizing antibodies is much lower (33 out of 101 tested). On the contrary, antibodies binding to the left-side (among which sotrovimab, bamlanivimab and cilgavimab), although their epitopes do not overlap with the ACE2 interaction region, are mostly neutralizing (21 out of 37 tested). As already discussed for sotrovimab, antibodies binding to this region impair the binding of the spike to the ACE2 dimer by their steric clashes with the second monomer (Supp. figure 1). A significant proportion of the antibodies binding to the vamp region is also neutralizing, although this region does not overlap with the ACE2 region. As antibodies binding to this region do not compete with most of the antibodies binding to the other regions, they might represent an interesting opportunity for combination treatments.

**Table 1:** Binding regions on the RBD. ^a^: secondary structures, as defined in **Erreur ! Source du renvoi introuvable.**. ^b^: solvent accessibility of the region in the opened and closed conformations. ^c^: number of antibodies of the set binding to this region. ^d^: list of the clusters binding to this region. ^e^: number of antibodies binding to CoV-2/ number of antibodies for which binding to CoV-2 has been tested. ^f^: number of antibodies binding to CoV-1/ number of antibodies for which binding to CoV-1 has been tested. ^g^: number of antibodies neutralizing CoV-2/ number of antibodies for which neutralization has been tested. ^h^: representatives binding to this region.

| Epitope | Sec. Struct.<sup>a</sup> | Buried closed/open<sup>b</sup> | Tot<sup>c</sup> | Clusters<sup>d</sup> | CoV2 B/tested<sup>e</sup> | CoV1 N/tested<sup>f</sup> | CoV2 neut/tested<sup>g</sup> | Antibodies<sup>h</sup> |
| --- | --- | --- | --- | --- | --- | --- | --- | --- |
| sole | L7S7L8 | no/no | 125 | 1; 26 | 112/112 | 35/84 | 33/101 | MR17, SR4, BD23, Fab2-4, CV07-250, S2M11, C002, C104, C121, C144, Nb6, COVA2-39, COVA2-04, P2C-1A3 |
| *right-sole* | H4L3H5S6<br>L6L7S7L8 | partial/no | 204 | 2; 5; 7;<br>9; 11;<br>14; 18 | 179/179 | 48/128 | 95/159 | **etevesimab**, casirivimab, CB6, B38, BD-604, BD-629, BD-236, 7CJF, CC12.1, CC12.3, C105, CV30, S2E12, C102, Fab-298, P2C-1F11, Fab2-17, FC05, COV2-2029 |
| *left-sole* | L5L6L7 | no/no | 61 | 6; 8 | 56/56 | 17/49 | 19/53 | **tixagevimab**, TY1, Sb23, BD-368-2, CV07-270, H11-D4, H11-H4, S2H13, C110, C119, P2B-2F6, S309 |
| *heel* | L1L5L8 | no/no | 1 |  | 1/1 | 0 | 1/1 | **imdevimab** |
| *left-side* | H1L1S1S9 | partial/partial | 48 | 10; 23;<br>24; 32 | 46/46 | 7/26 | 21/37 | **sotrovimab**, bamlanivimab, Fab-52, COV2-2346, 2M-9F10, COV2-2304, cilgavimab, C135, S2H14, m338 |
| *right-side* | H2L2S3H3<br>L3H4 | yes/no | 26 | 19; 21 | 26/26 | 16/24 | 13/26 | **VHH-72**, CR3022, EY6A, S2A4, S304, COV1-16, H014 |
| *rear* | H1S2H2L2<br>L5 | yes/no | 24 | 16; 31 | 23/23 | 2/18 | 18/22 | **COV2-2064**, regdanvimab, **C7** |
| *vamp* | H5L4L6L7 | yes/no | 121 | 3; 4; 12;<br>13; 15;<br>17; 20;<br>22; 25;<br>27; 28;<br>29; 30 | 191/192 | 94/174 | 37/127 | **CC12.7**, mAb-81, C021, COV2-2313, COV2-2589, COV2-2944, COV2-2821, COV2-2619, COV2-2619, CV07-200, COV2-2473, COV2-2022, COV2-2700, COV2-2449 |

This data can be used to predict non-competing antibodies, for example for combination therapies. Three such combinations are already in trial (tixagevimab + cilgavimab) or already authorized (etevesimab + bamlanivimab and casirivimab + imdevimab). In each case, the two combined antibodies can bind simultaneously to the RBD domain. Many other combinations of two non-competing neutralizing antibodies are possible, and easy to find using the presented data. Other combinations could also be tested, for example the Nanobody VHH-72, which is neutralizing^14^, does not compete with any of the antibodies under trial. Consequently, combination of VHH-72 with any of the 8 antibodies under trial could potentially increase their efficiency. VHH-72 could even be added to the combinations that have already proven efficiency.

## Variants of the virus

Binding of an antibody to its target can be affected by mutations of this target, especially within the epitope. SARS-CoV-2, as all viruses, is subjected to mutations, and a study as already shown that some of the naturally occurring mutations, but also other mutations that could appear, affect the neutralization efficiency of 3 of the antibodies under trial^15^. Epitope data that we have generated can be used to predict *in silico* the antibodies which might have their functional activity affected by existing mutations in the RBD domain of the spike protein. To this aim, known mutations were collected from the GISAID database^16^. A given mutation is predicted to have an impact on the binding of the antibodies belonging to a given group if it arises at a position belonging to the epitope of at least one antibody of the group. The predicted impact is also dependent on the nature of the mutation (table S2). A large number of mutations in the RBD have been found. A first evaluation of their impact on the binding of the different antibodies, but also on the binding to ACE2, can be obtained from this table.

The mutation N501Y, which is found in the new variant identified in Great-Britain, might affect efficiency of antibodies binding in the right-sole, sole or heel regions. It should be noted that this mutation has occurred multiple times, independently, around the world^‡^, indicating that it certainly brings advantages for the virus, such has the suspected higher contagiousness reported in the UK^§^. In the structure of the complex between imdevimab, casirivimab and RBD (PDB: 6XDG^11^), the side-chain of amino-acid N501 does not interact with the imdevimab, likely preserving the efficiency of the antibody (Supp. figure 3). Side-chain of N501 does not interact with casirivimab either; however, tyrosine being larger than glutamine, the N501Y mutation might introduce a supplementary interaction, which could reinforce the binding. In the predicted structure of the complex between RBD and etesevimab, N501 interacts with F27 (CDRH1). Thus, as for the interaction with ACE2, mutation N501Y could reinforce the interaction. Although effects of mutation are very difficult to predict, there is a good chance that in the present case the N501Y mutation could reinforce binding of RBD to ACE2, as well as to casirivimab and etesevimab.

In conclusion, AI-based methods allow to characterize sets of antibodies in a few hours, completely *in silico*, and compare them with other available antibodies. MAbCluster and MAbTope, the two methods used here are complementary. MAbCluster allows to group antibodies knowing only their sequences. Within one group, we demonstrated that all antibodies have overlapping epitopes. It is also possible to map biochemical and functional properties to the clustering, which allows to transfer the properties to antibodies of the same cluster for which these properties are unknown, with a high probability. MAbCluster can be applied even when the target’s 3D structure is unknown and cannot be accurately modeled from the structure of a close homolog. MAbCluster is very fast, and can be applied to millions of antibodies in a few hours.

When the 3D structure of the target is known, or can be modelled by homology, MAbTope allows to bring the information of the epitope for each cluster. MAbTope can be applied to thousands of antibodies in a few hours. This allows to rapidly choose promising candidates, predicted as having interesting structural, biochemical and functional properties. This also allows to select antibodies binding to “good” epitopes, those which can be predicted as important for the desired functional effect. Finally, new antibodies can be compared with existing ones, which can be important in terms of intellectual property, and potentially interesting combinations of two or more antibodies can be chosen.

## Acknowledgements

PRC and EA7501 were funded with support from the French National Research Agency under the program “Investissements d’avenir” Grant Agreement LabEx MabImprove: ANR-10-LABX-53; “ARD2020 Biomédicaments” projects GPCRAb, GPCRAb2 and BIO-S and “APR-IR MABSILICO” grants from Région Centre Val de Loire.

## Authors contributions

C.D. and A.M collected the data. Y.J., T.B. and A.P. developed the methods. Ph.R. contributed to the data visualizations. N.P., H.W. contributed to the discussion. A.P. analyzed the data. All authors contributed to the writing fo the paper.

## Material and methods

Antibody sequences targeting the spike protein of the different coronaviruses were collected from the database CoV-AbDab^17^, patent and literature mining and IMGT^18^. Clustering is made using the similarity measure defined in Bourquard et al^13^, then gathering groups of antibodies having pairwise similarity higher than 30. Cluster map is made using UMAP^19^. Epitope mapping is done using the method MAbTope^9^. Visualizations were made using Pymol^**^ for 3D structures and Observable^††^ for maps and matrices.

## Footnotes

* https://covid-nma.com/dataviz/

† https://clinicaltrials.gov/

‡ https://twitter.com/firefoxx66/status/1340359989395861506

§ https://www.ecdc.europa.eu/sites/default/files/documents/SARS-CoV-2-variant-multiple-spike-protein-mutations-United-Kingdom.pdf

** The PyMOL Molecular Graphics System, Version 2.0 Schrödinger, LLC.

†† https://observablehq.com/

