## Supplementary material for "Exploring epitope and functional diversity of anti-SARS-CoV2 antibodies using AI-based methods": Supp.

### Supplementary data

| Company | Antibody name | Trial phase | Seq | 3D |
|---|---|---|---|---|
| Junshi Biosciences/ Eli Lilly and Company/ AbCellera | etesevimab | Phase 2 | Yes | No |
| | bamlanivimab | Approved | Yes | No |
| | etesevimab+ bamlanivimab | Phase 2 | | |
| Tychan Pte. Ltd | TY027 | Phase 3 | No | No |
| Brii Biosciences | BRII-196 | Phase 1 | No | No |
| | BRII-198 | Phase 1 | No | No |
| AbbVie | ABBV-47D11 | Phase 1 | No | No |
| Sorrento Therapeutics, Inc. | COVI-GUARD | Phase 1 | No | No |
| | COVI-AMG | Phase 1/2 | No | No |
| Mabwell (Shanghai) Bioscience Co., Ltd. | MW33 | Phase 1 | No | No |
| HiFiBiO Therapeutics | HFB30132A | Phase 1 | No | No |
| Ology Bioservices | ADM03820 | Phase 1 | No | No |
| Hengenix Biotech Inc | HLX70 | Phase 1 | No | No |
| U. Cologne / Boehringer Ingelheim | DZIF-10c | Phase 1/2 | No | No |
| Beigene | BGB DXP593 | Phase 2 | No | No |
| Sinocelltech Ltd. | SCTA01 | Phase 2/3 | No | No |
| AstraZeneca | cilgavimab | Phase 1 | Yes | No |
| | tixagevimab | Phase 1 | Yes | No |
| | cilgavimab+ tixagevimab | Phase 3 | | |
| Celltrion | regdanvimab | Phase 2/3 | Yes | No |
| Vir Biotechnol./ GlaxoSmithKline | sotrovimab | Phase 2/3 | Yes | No* |
| Regeneron | casirivimab | Approved | Yes | Yes |
| | imdevimab | | Yes | Yes |
| | casirivimab+ imdevimab | Phase 3 | | |

*Supp. table 1: list of the anti-SARS-CoV-2 spike antibodies that have entered trials. *: antibody S309, for which the structure of the complex with the spike protein is available, has only one amino acid difference with sotrovimab.*

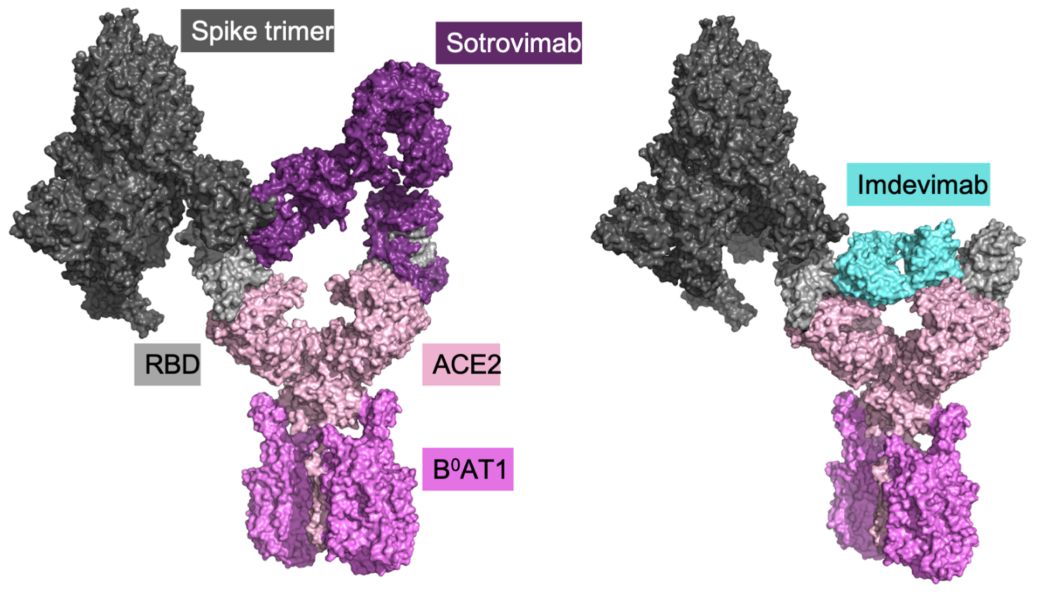


*Supp. figure 1: Sotrovimab (left) and imdevimab (right) are neutralizing because of steric hindrances with the second ACE2 monomer.*

| Cluster | Nb Abs | Representative |
|---|---|---|
| **1** | 94 | **C135**, **C002**, **imdevimab** |
| **2** | 75 | **P2C-1F11**, **B38**, **CV30**, **CC12.3**, **CC12.1**, **C102**, **COVA2-04**, **BD-629**, **BD-604**, **BD-236**, **7CJF** |
| **3** | 56 | mAb-81 |
| **4** | 43 | C021 |
| **5** | 36 | Fab2-17 |
| **6** | 24 | **C110** |
| **7** | 23 | COV2-2029 |
| **8** | 23 | **C119** |
| **9** | 22 | **Fab-298** |
| **10** | 18 | m338 |
| **11** | 16 | **S2E12** |
| **12** | 15 | COV2-2313 |
| **13** | 15 | COV2-2589 |
| **14** | 15 | etesevimab |
| **15** | 15 | COV2-2944 |
| **16** | 14 | COV2-2064 |
| **17** | 14 | COV2-2821 |
| **18** | 13 | FC05 |
| **19** | 13 | **S304** |
| **20** | 11 | COV2-2619 |
| **21** | 10 | **S2A4** |
| **22** | 10 | CV07-200 |
| **23** | 9 | COV2-2304 |
| **24** | 9 | COV2-2346 |
| **25** | 9 | COV2-2473 |
| **26** | 9 | **BD23** |
| **27** | 9 | COV2-2022 |
| **28** | 8 | COV2-2700 |
| **29** | 8 | COV2-2449 |
| **30** | 8 | CC12.7 |
| **31** | 7 | C7 |
| **32** | 7 | 2M-9F10 |
| **33** | 7 | VHH-21 |
| **34** | 2 | **S309**, sotrovimab |
| **35** | 1 | **Fab-52** |
| **36** | 3 | **S2H14** |
| **37** | 2 | **S2H13** |
| **38** | 2 | **casirivimab** |
| **39** | 1 | **CV07-250** |
| **40** | 4 | **CV07-270** |

| | | |
|---|---|---|
| **41** | 1 | **H014** |
| **42** | 1 | **BD-368-2** |
| **43** | 1 | **P2B-2F6** |
| **44** | 5 | **P2C-1A3** |
| **45** | 2 | **Fab2-4** |
| **46** | 1 | **COVA1-16** |
| **47** | 2 | **COVA2-39** |
| **48** | 1 | **C104** |
| **49** | 2 | **C105** |
| **50** | 5 | **C144** |
| **51** | 1 | **Nb#6** |
| **52** | 1 | **Sb23** |
| **53** | 1 | **SR4** |
| **54** | 3 | **MR17** |
| **55** | 1 | **VHH-72** |
| **56** | 2 | **H11-D4**, **H11-H4** |
| **57** | 1 | **S2M11** |

*Supp. table 2: list of antibodies clusters included in the study shown figure 4. Antibodies for which the 3D structure of the antibody-target complex is available are shown in bold.*

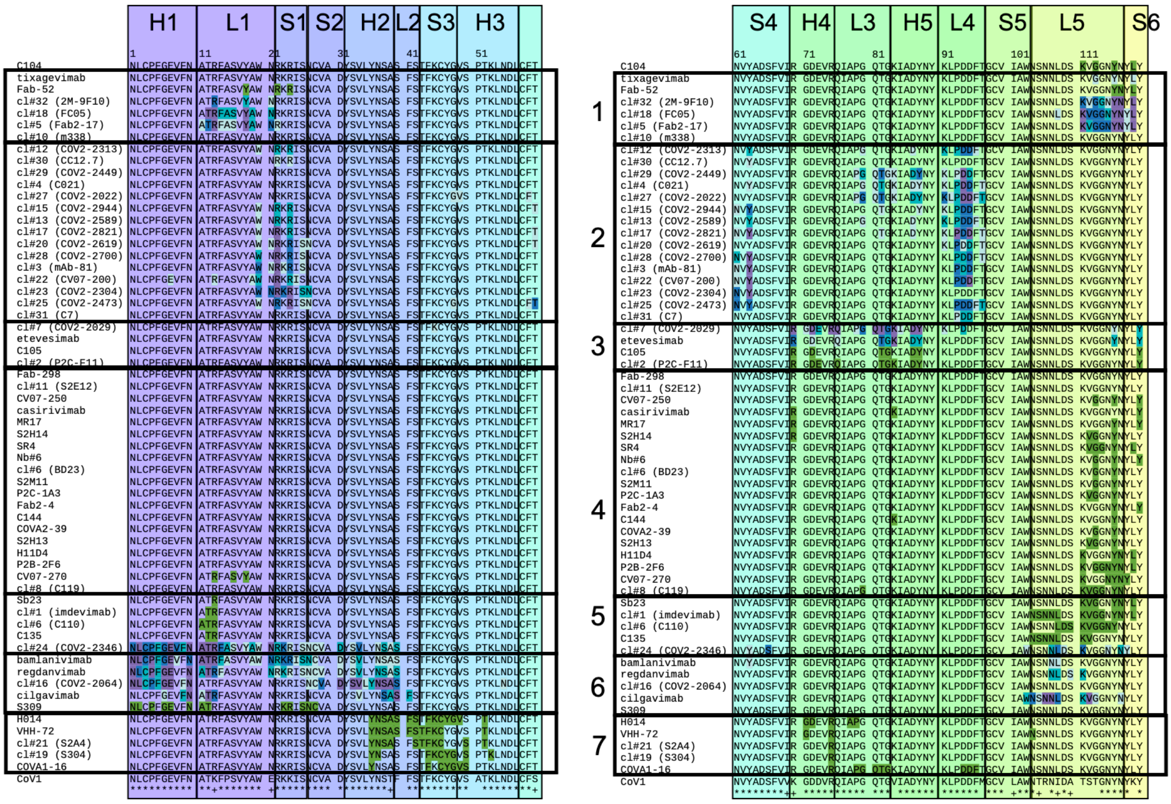


*Supp. figure 2: Epitopes of the representatives of each cluster. Epitopes deduced from crystal structures are indicated in green, epitopes predicted by MAbTope are indicated in blue depending on the probability (violet: very high, blue: high, cyan: medium, light cyan: low). The groups correspond to those indicated on the matrix Figure 4. The colored regions correspond to those indicated on Figure 2. The sequence of the CoV-1 RBD is indicated, with conserved (*) and similar (+) amino acids between CoV-2 and CoV-1 sequences.*

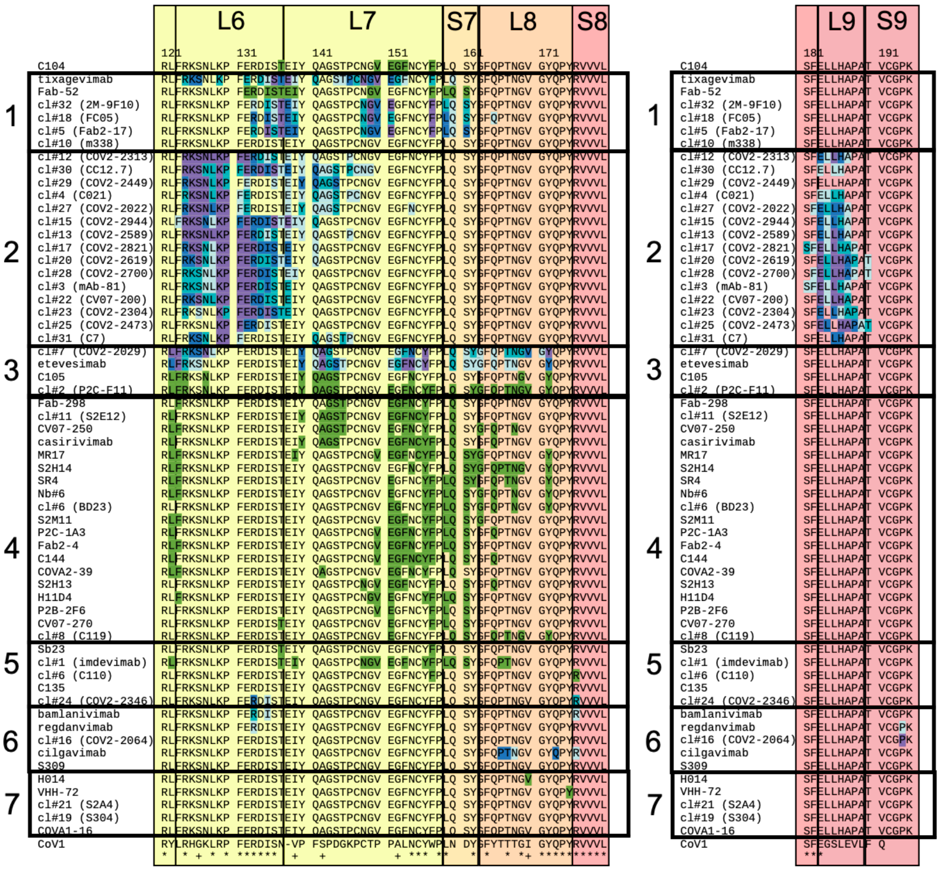


*Supp. Figure 2 (continued)*

| Name | C104 | grp1 | grp2 | grp3 | grp4 | grp5 | grp6 | grp7 | ACE2 | Ali. | pos | ref | alt | Nb. seq | Frequency (%) |
|---|---|---|---|---|---|---|---|---|---|---|---|---|---|---|---|
| Reference | | | | | | | | | | | | | | 47009 | 0,98263 |
| \|338\|F\|L | no | no | no | no | no | yes | yes | no | no | 5 | 338 | F | L | 1 | 2,1E-05 |
| \|339\|G\|D | no | no | no | no | no | yes | yes | no | no | 6 | 339 | G | D | 5 | 0,0001 |
| \|340\|E\|K | no | no | no | no | no | yes | yes | no | no | 7 | 340 | E | K | 5 | 0,0001 |
| \|341\|V\|I | no | no | no | no | no | yes | yes | no | no | 8 | 341 | V | I | 4 | 8,4E-05 |
| \|344\|A\|S | no | no | no | no | no | yes | yes | no | no | 11 | 344 | A | S | 32 | 0,00067 |
| \|344\|A\|T | no | no | no | no | no | yes | yes | no | no | 11 | 344 | A | T | 1 | 2,1E-05 |
| \|346\|R\|K | no | yes | no | no | no | yes | yes | no | no | 13 | 346 | R | K | 4 | 8,4E-05 |
| \|346\|R\|S | no | yes | no | no | no | yes | yes | no | no | 13 | 346 | R | S | 1 | 2,1E-05 |
| \|346\|R\|T | no | yes | no | no | no | yes | yes | no | no | 13 | 346 | R | T | 1 | 2,1E-05 |
| \|348\|A\|T | no | yes | no | no | no | yes | no | no | no | 15 | 348 | A | T | 1 | 2,1E-05 |
| \|348\|A\|S | no | yes | no | no | no | yes | no | no | no | 15 | 348 | A | S | 2 | 4,2E-05 |
| \|348\|A\|V | no | yes | no | no | no | yes | no | no | no | 15 | 348 | A | V | 1 | 2,1E-05 |
| \|352\|A\|S | no | yes | no | no | no | yes | no | no | no | 19 | 352 | A | S | 7 | 0,00015 |
| \|354\|N\|K | no | yes | yes | no | no | yes | yes | no | no | 21 | 354 | N | K | 5 | 0,0001 |
| \|354\|N\|D | no | yes | yes | no | no | yes | yes | no | no | 21 | 354 | N | D | 10 | 0,00021 |
| \|356\|K\|R | no | yes | yes | no | no | yes | yes | no | no | 23 | 356 | K | R | 5 | 0,0001 |

<table>
<tr><td>|357|R|K</td><td>no</td><td>yes</td><td>yes</td><td>no</td><td>no</td><td>yes</td><td>yes</td><td>no</td><td>no</td><td>24</td><td>357</td><td>R</td><td>K</td><td>6</td><td>0,00013</td></tr>
<tr><td>|359|S|N</td><td>no</td><td>yes</td><td>yes</td><td>no</td><td>no</td><td>yes</td><td>yes</td><td>no</td><td>no</td><td>26</td><td>359</td><td>S</td><td>N</td><td>1</td><td>2,1E-05</td></tr>
<tr><td>|361|C|S</td><td>no</td><td>yes</td><td>yes</td><td>no</td><td>no</td><td>yes</td><td>yes</td><td>no</td><td>no</td><td>28</td><td>361</td><td>C</td><td>S</td><td>4</td><td>8,4E-05</td></tr>
<tr><td>|361|C|_</td><td>no</td><td>yes</td><td>yes</td><td>no</td><td>no</td><td>yes</td><td>yes</td><td>no</td><td>no</td><td>28</td><td>361</td><td>C</td><td>_</td><td>2</td><td>4,2E-05</td></tr>
<tr><td>|362|V|F</td><td>no</td><td>yes</td><td>yes</td><td>no</td><td>no</td><td>yes</td><td>yes</td><td>no</td><td>no</td><td>29</td><td>362</td><td>V</td><td>F</td><td>3</td><td>6,3E-05</td></tr>
<tr><td>|365|Y|S</td><td>no</td><td>yes</td><td>yes</td><td>no</td><td>no</td><td>yes</td><td>yes</td><td>no</td><td>no</td><td>32</td><td>365</td><td>Y</td><td>S</td><td>3</td><td>6,3E-05</td></tr>
<tr><td>|365|Y|D</td><td>no</td><td>yes</td><td>yes</td><td>no</td><td>no</td><td>yes</td><td>yes</td><td>no</td><td>no</td><td>32</td><td>365</td><td>Y</td><td>D</td><td>5</td><td>0,0001</td></tr>
<tr><td>|367|V|F</td><td>no</td><td>yes</td><td>yes</td><td>no</td><td>no</td><td>yes</td><td>yes</td><td>no</td><td>no</td><td>34</td><td>367</td><td>V</td><td>F</td><td>27</td><td>0,00056</td></tr>
<tr><td>|367|V|I</td><td>no</td><td>yes</td><td>yes</td><td>no</td><td>no</td><td>yes</td><td>yes</td><td>no</td><td>no</td><td>34</td><td>367</td><td>V</td><td>I</td><td>4</td><td>8,4E-05</td></tr>
<tr><td>|370|N|K</td><td>no</td><td>no</td><td>no</td><td>no</td><td>no</td><td>yes</td><td>yes</td><td>yes</td><td>no</td><td>37</td><td>370</td><td>N</td><td>K</td><td>3</td><td>6,3E-05</td></tr>
<tr><td>|370|N|S</td><td>no</td><td>no</td><td>no</td><td>no</td><td>no</td><td>yes</td><td>yes</td><td>yes</td><td>no</td><td>37</td><td>370</td><td>N</td><td>S</td><td>1</td><td>2,1E-05</td></tr>
<tr><td>|371|S|P</td><td>no</td><td>no</td><td>no</td><td>no</td><td>no</td><td>yes</td><td>yes</td><td>yes</td><td>no</td><td>38</td><td>371</td><td>S</td><td>P</td><td>2</td><td>4,2E-05</td></tr>
<tr><td>|371|S|T</td><td>no</td><td>no</td><td>no</td><td>no</td><td>no</td><td>yes</td><td>yes</td><td>yes</td><td>no</td><td>38</td><td>371</td><td>S</td><td>T</td><td>3</td><td>6,3E-05</td></tr>
<tr><td>|373|S|L</td><td>no</td><td>no</td><td>no</td><td>no</td><td>no</td><td>yes</td><td>yes</td><td>yes</td><td>no</td><td>40</td><td>373</td><td>S</td><td>L</td><td>4</td><td>8,4E-05</td></tr>
<tr><td>|376|T|I</td><td>no</td><td>no</td><td>no</td><td>no</td><td>no</td><td>no</td><td>no</td><td>yes</td><td>no</td><td>43</td><td>376</td><td>T</td><td>I</td><td>6</td><td>0,00013</td></tr>
<tr><td>|382|V|L</td><td>no</td><td>no</td><td>no</td><td>no</td><td>no</td><td>no</td><td>no</td><td>yes</td><td>no</td><td>49</td><td>382</td><td>V</td><td>L</td><td>40</td><td>0,00084</td></tr>
<tr><td>|384|P|S</td><td>no</td><td>no</td><td>no</td><td>no</td><td>no</td><td>no</td><td>no</td><td>yes</td><td>no</td><td>51</td><td>384</td><td>P</td><td>S</td><td>5</td><td>0,0001</td></tr>
<tr><td>|384|P|L</td><td>no</td><td>no</td><td>no</td><td>no</td><td>no</td><td>no</td><td>no</td><td>yes</td><td>no</td><td>51</td><td>384</td><td>P</td><td>L</td><td>11</td><td>0,00023</td></tr>
<tr><td>|384|P|R</td><td>no</td><td>no</td><td>no</td><td>no</td><td>no</td><td>no</td><td>no</td><td>yes</td><td>no</td><td>51</td><td>384</td><td>P</td><td>R</td><td>13</td><td>0,00027</td></tr>
<tr><td>|385|T|I</td><td>no</td><td>no</td><td>no</td><td>no</td><td>no</td><td>no</td><td>no</td><td>yes</td><td>no</td><td>52</td><td>385</td><td>T</td><td>I</td><td>3</td><td>6,3E-05</td></tr>
<tr><td>|401|V|L</td><td>no</td><td>no</td><td>no</td><td>no</td><td>no</td><td>no</td><td>no</td><td>no</td><td>no</td><td>68</td><td>401</td><td>V</td><td>L</td><td>2</td><td>4,2E-05</td></tr>
<tr><td>|402|I|V</td><td>no</td><td>no</td><td>no</td><td>no</td><td>no</td><td>no</td><td>no</td><td>no</td><td>no</td><td>69</td><td>402</td><td>I</td><td>V</td><td>4</td><td>8,4E-05</td></tr>
<tr><td>|403|R|K</td><td>no</td><td>no</td><td>no</td><td>no</td><td>no</td><td>no</td><td>no</td><td>no</td><td>no</td><td>70</td><td>403</td><td>R</td><td>K</td><td>11</td><td>0,00023</td></tr>
<tr><td>|406|E|Q</td><td>no</td><td>no</td><td>yes</td><td>no</td><td>no</td><td>no</td><td>no</td><td>no</td><td>no</td><td>73</td><td>406</td><td>E</td><td>Q</td><td>7</td><td>0,00015</td></tr>
<tr><td>|408|R|K</td><td>no</td><td>no</td><td>yes</td><td>no</td><td>no</td><td>no</td><td>no</td><td>yes</td><td>no</td><td>75</td><td>408</td><td>R</td><td>K</td><td>1</td><td>2,1E-05</td></tr>
<tr><td>|408|R|T</td><td>no</td><td>no</td><td>yes</td><td>no</td><td>no</td><td>no</td><td>no</td><td>yes</td><td>no</td><td>75</td><td>408</td><td>R</td><td>T</td><td>4</td><td>8,4E-05</td></tr>
<tr><td>|408|R|I</td><td>no</td><td>no</td><td>yes</td><td>no</td><td>no</td><td>no</td><td>no</td><td>yes</td><td>no</td><td>75</td><td>408</td><td>R</td><td>I</td><td>1</td><td>2,1E-05</td></tr>
<tr><td>|410|I|V</td><td>no</td><td>no</td><td>no</td><td>no</td><td>no</td><td>no</td><td>no</td><td>yes</td><td>no</td><td>77</td><td>410</td><td>I</td><td>V</td><td>1</td><td>2,1E-05</td></tr>
<tr><td>|411|A|S</td><td>no</td><td>no</td><td>no</td><td>no</td><td>no</td><td>no</td><td>no</td><td>yes</td><td>no</td><td>78</td><td>411</td><td>A</td><td>S</td><td>9</td><td>0,00019</td></tr>
<tr><td>|414|Q|R</td><td>no</td><td>no</td><td>yes</td><td>yes</td><td>no</td><td>no</td><td>no</td><td>yes</td><td>no</td><td>81</td><td>414</td><td>Q</td><td>R</td><td>1</td><td>2,1E-05</td></tr>
<tr><td>|427|D|Y</td><td>no</td><td>no</td><td>yes</td><td>yes</td><td>no</td><td>no</td><td>no</td><td>yes</td><td>no</td><td>94</td><td>427</td><td>D</td><td>Y</td><td>1</td><td>2,1E-05</td></tr>
<tr><td>|430|T|I</td><td>no</td><td>no</td><td>yes</td><td>no</td><td>no</td><td>no</td><td>no</td><td>no</td><td>no</td><td>97</td><td>430</td><td>T</td><td>I</td><td>3</td><td>6,3E-05</td></tr>
<tr><td>|439|N|K</td><td>no</td><td>no</td><td>no</td><td>no</td><td>no</td><td>yes</td><td>yes</td><td>no</td><td>no</td><td>106</td><td>439</td><td>N</td><td>K</td><td>3</td><td>6,3E-05</td></tr>
<tr><td>|444|K|N</td><td>yes</td><td>yes</td><td>no</td><td>no</td><td>yes</td><td>yes</td><td>yes</td><td>no</td><td>no</td><td>111</td><td>444</td><td>K</td><td>N</td><td>3</td><td>6,3E-05</td></tr>
<tr><td>|446|G|V</td><td>yes</td><td>yes</td><td>no</td><td>no</td><td>yes</td><td>yes</td><td>yes</td><td>no</td><td>yes</td><td>113</td><td>446</td><td>G</td><td>V</td><td>7</td><td>0,00015</td></tr>
<tr><td>|450|N|K</td><td>yes</td><td>yes</td><td>no</td><td>yes</td><td>yes</td><td>yes</td><td>yes</td><td>no</td><td>no</td><td>117</td><td>450</td><td>N</td><td>K</td><td>3</td><td>6,3E-05</td></tr>
<tr><td>|452|L|R</td><td>yes</td><td>yes</td><td>no</td><td>yes</td><td>yes</td><td>yes</td><td>no</td><td>no</td><td>no</td><td>119</td><td>452</td><td>L</td><td>R</td><td>12</td><td>0,00025</td></tr>
<tr><td>|453|Y|F</td><td>no</td><td>no</td><td>no</td><td>yes</td><td>yes</td><td>no</td><td>no</td><td>no</td><td>yes</td><td>120</td><td>453</td><td>Y</td><td>F</td><td>18</td><td>0,00038</td></tr>
<tr><td>|455|L|F</td><td>no</td><td>no</td><td>no</td><td>yes</td><td>yes</td><td>yes</td><td>no</td><td>no</td><td>yes</td><td>122</td><td>455</td><td>L</td><td>F</td><td>3</td><td>6,3E-05</td></tr>
<tr><td>|456|F|L</td><td>no</td><td>no</td><td>yes</td><td>yes</td><td>yes</td><td>no</td><td>no</td><td>no</td><td>yes</td><td>123</td><td>456</td><td>F</td><td>L</td><td>2</td><td>4,2E-05</td></tr>
<tr><td>|463|P|S</td><td>no</td><td>yes</td><td>yes</td><td>no</td><td>no</td><td>no</td><td>no</td><td>no</td><td>no</td><td>130</td><td>463</td><td>P</td><td>S</td><td>6</td><td>0,00013</td></tr>
<tr><td>|475|A|V</td><td>no</td><td>no</td><td>yes</td><td>yes</td><td>yes</td><td>no</td><td>no</td><td>no</td><td>yes</td><td>142</td><td>475</td><td>A</td><td>V</td><td>10</td><td>0,00021</td></tr>
</table>

| | | | | | | | | | | | | | | |
|---|---|---|---|---|---|---|---|---|---|---|---|---|---|---|
| \|476\|G\|S | no | no | yes | yes | yes | no | no | no | yes | 143 | 476 | G | S | 13 | 0,00027 |
| \|477\|S\|N | no | yes | yes | yes | yes | no | no | no | yes | 144 | 477 | S | N | 15 | 0,00031 |
| \|477\|S\|I | no | yes | yes | yes | yes | no | no | no | yes | 144 | 477 | S | I | 15 | 0,00031 |
| \|478\|T\|K | no | yes | yes | yes | yes | no | no | no | no | 145 | 478 | T | K | 7 | 0,00015 |
| \|479\|P\|L | no | yes | yes | yes | yes | no | no | no | no | 146 | 479 | P | L | 2 | 4,2E-05 |
| \|479\|P\|S | no | yes | yes | yes | yes | no | no | no | no | 146 | 479 | P | S | 3 | 6,3E-05 |
| \|483\|V\|A | yes | no | no | no | yes | yes | no | no | no | 150 | 483 | V | A | 51 | 0,00107 |
| \|483\|V\|F | yes | no | no | no | yes | yes | no | no | no | 150 | 483 | V | F | 10 | 0,00021 |
| \|484\|E\|K | yes | yes | no | yes | yes | yes | no | no | no | 151 | 484 | E | K | 11 | 0,00023 |
| \|484\|E\|Q | yes | yes | no | yes | yes | yes | no | no | no | 151 | 484 | E | Q | 4 | 8,4E-05 |
| \|484\|E\|A | yes | yes | no | yes | yes | yes | no | no | no | 151 | 484 | E | A | 3 | 6,3E-05 |
| \|486\|F\|L | yes | yes | no | yes | yes | yes | no | no | yes | 153 | 486 | F | L | 2 | 4,2E-05 |
| \|490\|F\|L | yes | yes | no | yes | yes | yes | no | no | yes | 157 | 490 | F | L | 3 | 6,3E-05 |
| \|493\|Q\|L | no | yes | no | yes | yes | yes | no | no | yes | 160 | 493 | Q | L | 10 | 0,00021 |
| \|493\|Q\|K | no | yes | no | yes | yes | yes | no | no | yes | 160 | 493 | Q | K | 3 | 6,3E-05 |
| \|494\|S\|P | no | yes | no | yes | yes | yes | no | no | yes | 161 | 494 | S | P | 56 | 0,00117 |
| \|494\|S\|A | no | yes | no | yes | yes | yes | no | no | yes | 161 | 494 | S | A | 2 | 4,2E-05 |
| \|501\|N\|Y | no | no | no | yes | yes | no | yes | no | yes | 168 | 501 | N | Y | 8 | 0,00017 |
| \|501\|N\|T | no | no | no | yes | yes | no | yes | no | yes | 168 | 501 | N | T | 21 | 0,00044 |
| \|503\|V\|F | no | no | no | yes | no | no | no | yes | yes | 170 | 503 | V | F | 2 | 4,2E-05 |
| \|505\|Y\|H | no | no | no | yes | yes | no | no | no | yes | 172 | 505 | Y | H | 2 | 4,2E-05 |
| \|505\|Y\|C | no | no | no | yes | yes | no | no | no | yes | 172 | 505 | Y | C | 18 | 0,00038 |
| \|505\|Y\|_ | no | no | no | yes | yes | no | no | no | yes | 172 | 505 | Y | _ | 18 | 0,00038 |
| \|506\|Q\|K | no | no | no | no | no | no | yes | no | no | 173 | 506 | Q | K | 14 | 0,00029 |
| \|508\|Y\|H | no | no | no | no | no | no | no | yes | no | 175 | 508 | Y | H | 2 | 4,2E-05 |
| \|513\|L\|F | no | no | yes | no | no | no | no | no | no | 180 | 513 | L | F | 2 | 4,2E-05 |
| \|516\|E\|Q | no | no | yes | no | no | no | no | no | no | 183 | 516 | E | Q | 3 | 6,3E-05 |
| \|517\|L\|F | no | no | yes | no | no | no | no | no | no | 184 | 517 | L | F | 2 | 4,2E-05 |
| \|520\|A\|S | no | no | yes | no | no | no | no | no | no | 187 | 520 | A | S | 100 | 0,00209 |
| \|521\|P\|R | no | no | yes | no | no | no | no | no | no | 188 | 521 | P | R | 18 | 0,00038 |
| \|521\|P\|S | no | no | yes | no | no | no | no | no | no | 188 | 521 | P | S | 3 | 6,3E-05 |
| \|522\|A\|V | no | no | yes | no | no | no | no | no | no | 189 | 522 | A | V | 13 | 0,00027 |
| \|522\|A\|S | no | no | yes | no | no | no | no | no | no | 189 | 522 | A | S | 57 | 0,00119 |
| \|522\|A\|G | no | no | yes | no | no | no | no | no | no | 189 | 522 | A | G | 3 | 6,3E-05 |

*Supp. table 3: Known mutants of the RBD domain. Ali: position in the alignment of figure S1; pos: position in the complete spike sequence; ref: amino acid at this position in the reference sequence; alt: amino acid at this position in the mutated sequence; nb seq: number of sequenced samples in which this mutation was found. Name are colored depending on the nature of the mutation: yellow for mutations involving small modifications of physico-chemical properties, orange for medium modifications, red for important modifications. Columns C104, grp1, …, grp7 give an estimation of the perturbation involved by the mutation on the binding of antibodies belonging to each group. When a perturbation is predicted to arise, the cell is colored as a function of the importance of the physico-chemical properties modification.*

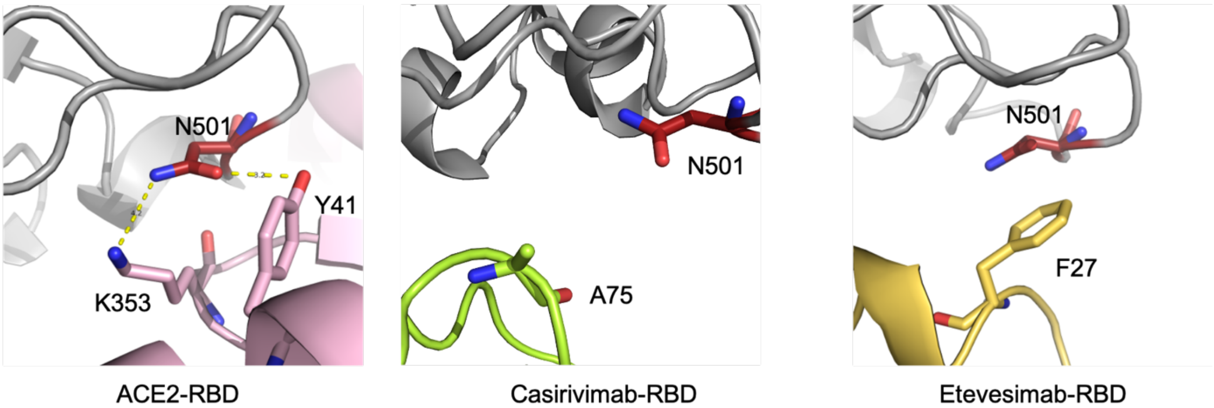


*Supp. figure 3: Interactions of amino acid N501 in complexes between RBD and ACE2 (left), casirivimab (center) and etevesimab (right).*
